# Modeling of mitochondrial genetic polymorphisms reveals induction of heteroplasmy by pleiotropic disease locus MT:10398A>G

**DOI:** 10.1101/2023.06.22.546129

**Authors:** Molly Smullen, Meagan N Olson, Liam F Murray, Madhusoodhanan Suresh, Guang Yan, Pepper Dawes, Nathaniel J Barton, Jivanna N Mason, Yucheng Zhang, Aria A Fernandez-Fontaine, George M Church, Diego Mastroeni, Qi Wang, Elaine T Lim, Yingleong Chan, Benjamin Readhead

**Author notes:** These authors contributed equally (Yingleong Chan, Benjamin Readhead). Corresponding authors Benjamin Readhead Yingleong Chan.

## Abstract

Mitochondrial (MT) dysfunction has been associated with several neurodegenerative diseases including Alzheimer’s disease (AD). While MT-copy number differences have been implicated in AD, the effect of MT heteroplasmy on AD has not been well characterized. Here, we analyzed over 1,800 whole genome sequencing data from four AD cohorts in seven different tissue types to determine the extent of MT heteroplasmy present. While MT-heteroplasmy was present throughout the entire MT genome for blood samples, we detected MT-heteroplasmy only within the MT control region for brain samples. We observed that an MT variant 10398A>G (rs2853826) was significantly associated with overall MT-heteroplasmy in brain tissue while also being linked with the largest number of distinct disease phenotypes of all annotated MT variants in *MitoMap*. Using gene-expression data from our brain samples, our modeling discovered several gene networks involved in mitochondrial respiratory chain and Complex I function associated with 10398A>G. The variant was also found to be an expression quantitative trait loci (eQTL) for the gene MT-ND3. We further characterized the effect of 10398A>G by phenotyping a population of lymphoblastoid cell-lines (LCLs) with and without the variant allele. Examination of RNA sequence data from these LCLs reveal that 10398A>G was an eQTL for MT-ND4. We also observed in LCLs that 10398A>G was significantly associated with overall MT-heteroplasmy within the MT control region, confirming the initial findings observed in post-mortem brain tissue. These results provide novel evidence linking MT SNPs with MT heteroplasmy and open novel avenues for the investigation of pathomechanisms that are driven by this pleiotropic disease associated loci.

## Introduction

Mutations in the MT genome have been reported to cause a wide variety of human diseases^1^. In particular, the oxidative phosphorylation (OXPHOS) function of the mitochondria has been shown to be disrupted by these mutations^2^, and defects in OXPHOS are linked with an increased risk of cancer, including prostate and breast cancer^3,4^. Besides mutations, dysregulation of mitochondria leading to metabolic defects have also been associated with Alzheimer’s disease (AD)^5,6^. By analyzing large metabolic data sets, researchers have shown that at later stages of AD, there is a change of energy utilization from fatty acids to amino acids and glucose, indicating an AD-associated switch in energy substrate utilization^7^. Great effort has been made to test the association of AD pathogenesis on a number of phenotypes related to mitochondrial dysfunction, such as overexpression of reactive oxygen species (ROS), calcium imbalance, and other defects of mitochondrial function and dynamics^8^. There is also evidence of significantly lower MT-DNA copy number in the temporal cortex^9^ and dorsolateral prefrontal cortex^10^ of AD patient brains compared to control subjects. Lastly, APOE ɛ4 is the common DNA variant that confers the highest increase in risk and age of onset of sporadic AD^11^ and was associated with impaired MT structure and function from proteins measured within postmortem human brain tissues of AD patients^12^. As such, studying the role of MT function in AD pathogenesis is currently an active area of research^13,14^.

Here, we primarily focus on MT heteroplasmy as a potential mechanism affecting MT function and thus impacting on risk of disease. MT heteroplasmy refers to the presence of MT mutations that occur in only a fraction of mitochondrial DNA within a given sample^15^. We analyzed data from 1,801 whole-genome sequenced (WGS) post-mortem tissue samples generated by the Accelerating Medicines Partnership in Alzheimer’s Disease (AMP-AD) for the presence of MT heteroplasmy. After discovering a significant association between the mitochondrial SNP 10398A>G (rs2853826) and MT heteroplasmy, we characterized the gene networks and gene expression changes associated with this single-nucleotide polymorphism (SNP). 10398A>G corresponds to the Thr114Ala missense mutation of the MT-ND3 gene coded by the MT, and is a common variant found within most populations with a reported allele frequency of 41.8% from the gnomAD database v3.1.1^16^. The 10398A>G allele has been collectively associated with an expansive set of phenotypically diverse diseases including AD^17^, Parkinson’s disease^18–24^, breast cancer^25–34^, gastric cancer, type 2 diabetes and several other human diseases and phenotypes^35–39^.

To explore the effect of 10398A>G on transcriptomic and MT variables, we performed experiments using lymphoblastoid cell lines (LCLs) from the Harvard Personal Genome Project (PGP)^40,41^. The PGP consists of many participants that have high coverage WGS data as well as self-reported phenotypic data that are publicly available for research^42,43^. We selected 60 participant LCLs that vary by the 10398 genotype (major allele A: n=30, minor allele G: n=30) and performed bulk RNA sequencing, as well as quantifying MT copy number and heteroplasmy, and report our findings herein.

## Results

### Significant heteroplasmy detected within the mitochondria control region from brain samples

We analyzed WGS data generated from 1,801 post-mortem tissue samples collected as part of the Religious Order Study (ROS), Memory and Aging Project (MAP), Mount Sinai Brain Bank (MSBB) and Mayo Temporal Cortex (MAYO) studies. Across these four studies, WGS data was available from a total of seven different tissues (**Table 1**). We analyzed data from these samples to perform MT variant calling and to estimate MT heteroplasmy at each MT genomic site (MT heteroplasmy was classified as present if at least 5% of reads that were reliably mapped to either the major or minor allele were discordant with the remaining reads).

**Table 1.** The various cohorts and samples used for analyzing whole-genome sequencing data for mitochondrial heteroplasmy. Samples were sequenced from the ROS (Religious Order Study), MAP (Memory and Aging Project), MSBB (Mount Sinai Brain Bank) and MAYO (Mayo Clinic) samples.

| Cohort | WGS Tissue | #Samples |
| --- | --- | --- |
| ROS | Blood | 227 |
|  | DLPFC (Dorsolateral prefrontal cortex) | 211 |
|  | PCC (Posterior cingulate cortex) | 38 |
|  | Cerebellum | 91 |
| MAP | Blood | 147 |
|  | DLPFC (Dorsolateral prefrontal cortex) | 240 |
|  | PCC (Posterior cingulate cortex) | 29 |
|  | Cerebellum | 155 |
| MSBB | BM-10 (Anterior prefrontal cortex) | 80 |
|  | BM-22 (Superior Temporal Gyrus) | 246 |
| MAYO | TCX (Temporal Cortex) | 337 |
| **Total** |  | **1801** |

While MT heteroplasmy was robustly detected in many samples, there was much higher overall heteroplasmy within the blood compared to the brain samples (**Figure 1A**). When the analysis was restricted to only brain samples, we detected significantly more heteroplasmy in the cortically derived samples than the cerebellum derived samples (**Figure 1B**).

**Figure 1.**
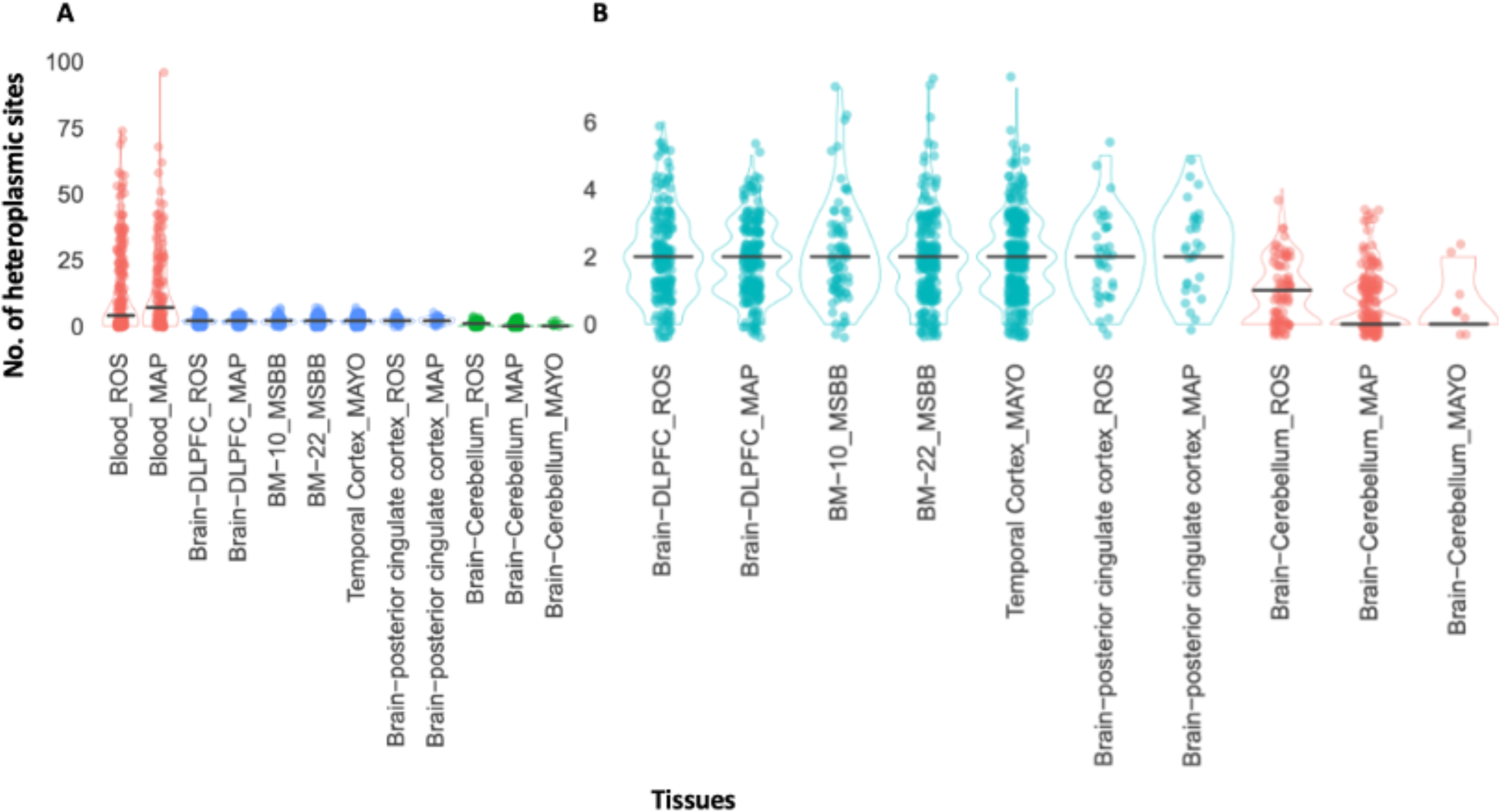
The number of heteroplasmic sites detected for each sample across the various tissues and cohorts. (A) Comparing the number of heteroplasmic sites in the blood and brain samples. (B) Comparing the number of heteroplasmic sites in only brain samples.

We observed that MT heteroplasmy detected within the blood derived samples demonstrated a distribution across the entire MT genome, whereas in the brain derived samples, heteroplasmy was entirely restricted to a set of several hundred bases corresponding to the MT control region (**Figure 2**), a highly polymorphic^44^, non-coding region of the MT genome which contains the origins of both transcription and replication^45^.

**Figure 2.**
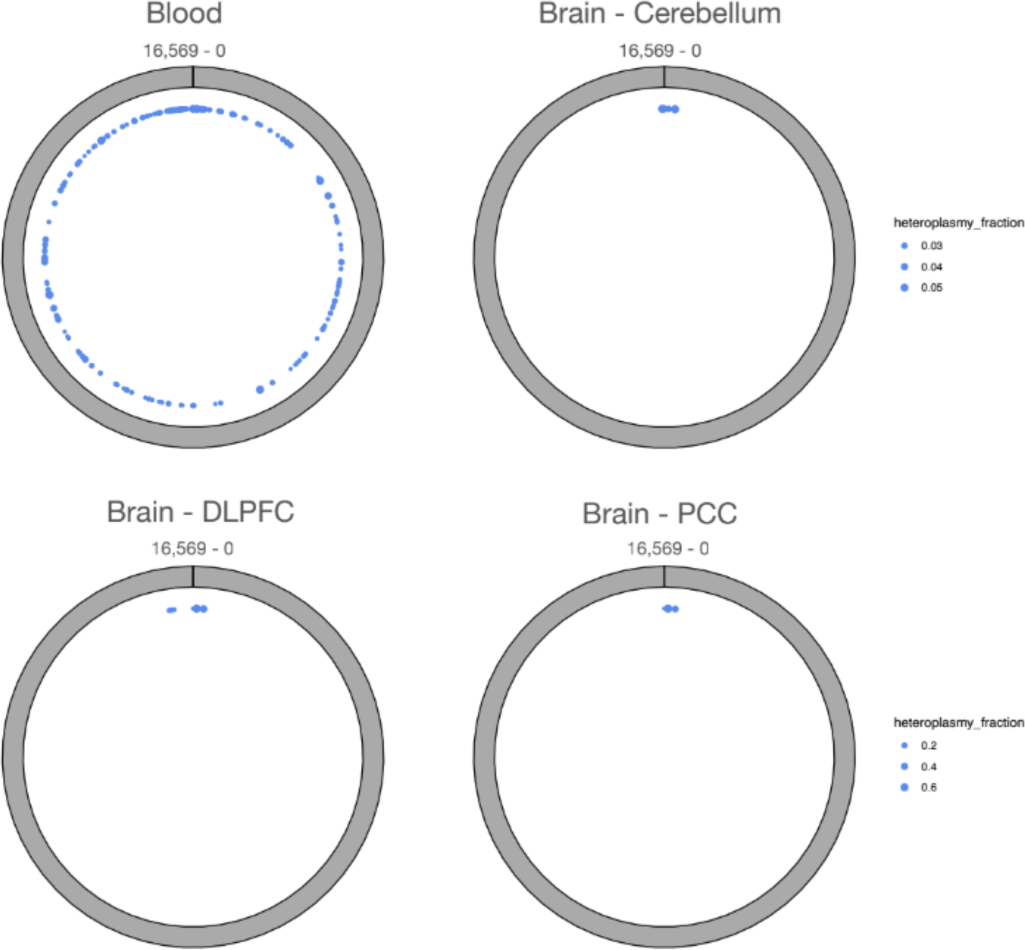
The location of the heteroplasmic sites detected for each tissue type across the mitochondrial genome.

### 10398A>G is significantly associated with mitochondrial heteroplasmy

Based on the analysis of mitochondrial heteroplasmy within brain samples, we performed a cis-heteroplasmy-QTL across the mitochondrial to determine if any SNPs were associated with heteroplasmy. We restricted this analysis to the brain regions with the largest subset of WGS samples, the dorsolateral prefrontal cortext (DLPFC) from the ROS / MAP studies (n=451). While a number of SNPs were found to be significantly associated with heteroplasmy, the 10398A>G variant had the most abundant signal, being significantly associated with heteroplasmy at five separate loci (FDR < 0.05) (**Table 2**). We further annotated each variant for its associated disease phenotypes using MITOMAP^46^ and observed that 10398A>G was associated with the largest number of distinct phenotypes of any reported MT variant (n=10). 10398A>G has been linked with AD^17^, Parkinson’s disease^18–24^, breast cancer^25–34^, type 2 diabetes and many other diseases and conditions^35–39^.

**Table 2.** Inherited heteroplasmy QTL alleles ranked according to number of significantly associated heteroplasmy sites (FDR < 0.05). Phenotype annotations from MITOMAP

| MT hetQTL locus | # Heteroplasmy sites | MT Heteroplasmy Sites | Estimates (Range) | FDR (Range) | # MITOMAP Diseases | MITOMAP Diseases |
| --- | --- | --- | --- | --- | --- | --- |
| 10398_A/G | 5 | 16129_G/A, 189_A/G, 185_G/A, 65_TG/T, 16093_T/C | 1.27-2.33 | 4.69e-02-1.35e-07 | 10 | PD protective factor / longevity / altered cell pH / metabolic syndrome / breast cancer risk / Leigh Syndrome risk / ADHD / cognitive decline / SCA2 age of onset / Fuchs endothelial corneal dystrophy |
| 11467_A/G | 4 | 65_TG/T, 189_A/G, 16093_T/C, 16188_CT/C | 1.15-3.22 | 4.72e-02-1.29e-06 | 2 | Altered brain pH / sCJD patients |
| 12308_A/G | 4 | 65_TG/T, 189_A/G, 16093_T/C, 16188_CT/C | 1.15-3.22 | 4.72e-02-1.29e-06 | 5 | CPEO / Stroke / CM / Breast & Renal & Prostate Cancer Risk / Altered brain pH / sCJD |
| 12372_G/A | 4 | 65_TG/T, 189_A/G, 16093_T/C, 16188_CT/C | 1.15-3.22 | 4.72e-02-1.29e-06 | 2 | Altered brain pH / sCJD patients |
| 14798_T/C | 4 | 185_G/A, 189_A/G, 65_TG/T, 16093_T/C | 1.22-2.79 | 1.45e-03-1.80e-06 |  |  |
| 11251_A/G | 3 | 185_G/A, 189_A/G, 65_TG/T | 1.31-2.61 | 1.07e-04-7E-06 | 1 | Reduced risk of PD |
| 11719_G/A | 3 | 189_A/G, 65_TG/T, 16129_G/A | 1.05-2.07 | 4.26e-02-1.08e-14 |  |  |
| 12612_A/G | 3 | 185_G/A, 189_A/G, 65_TG/T | 1.18-3.58 | 2.62e-02-2.89e-08 |  |  |
| 13617_T/C | 3 | 65_TG/T, 16188_CT/C, 185_G/A | 1.29-2.80 | 3.50e-02-8.59e-03 |  |  |
| 13708_G/A | 3 | 185_G/A, 189_A/G, 65_TG/T | 0.93-3.30 | 4.94e-02-1.58e-07 | 3 | LHON / Increased MS risk / higher freq in PD-ADS |

### The 10398A>G variant is significantly associated with MT-ND3 expression in human prefrontal cortex

Given the association of 10398A>G with mitochondrial heteroplasmy and diverse disease phenotypes, we analyzed its association with gene expression changes. By integrating RNA sequence data available on ROS / MAP cohort DLPFC samples, we performed a cis-eQTL analysis and found that 10398A>G was associated with MT-ND3 (Mitochondrially Encoded NADH Dehydrogenase 3) expression, where the minor G allele is associated with decreased expression (**Figure 3A**). MT-ND3 is one of the subunits of the mitochondrial respiratory chain complex I that enables NADH dehydrogenase (ubiquinone) activity. We thus constructed a targeted gene regulatory network aimed at identifying genes that are downstream of MT-ND3, conditioned on the relationship with 10398A>G, using a causal inference testing approach^47^ and identified a number of significantly associated downstream genes (**Figure 3B**). We then performed a gene set enrichment analysis on the MT-ND3 subnetwork using *Enrichr*^48–50^ and observed several enrichments for NADH dehydrogenase activity and Mitochondrial respiratory chain Complex I components (**Table 3**).

**Figure 3.**
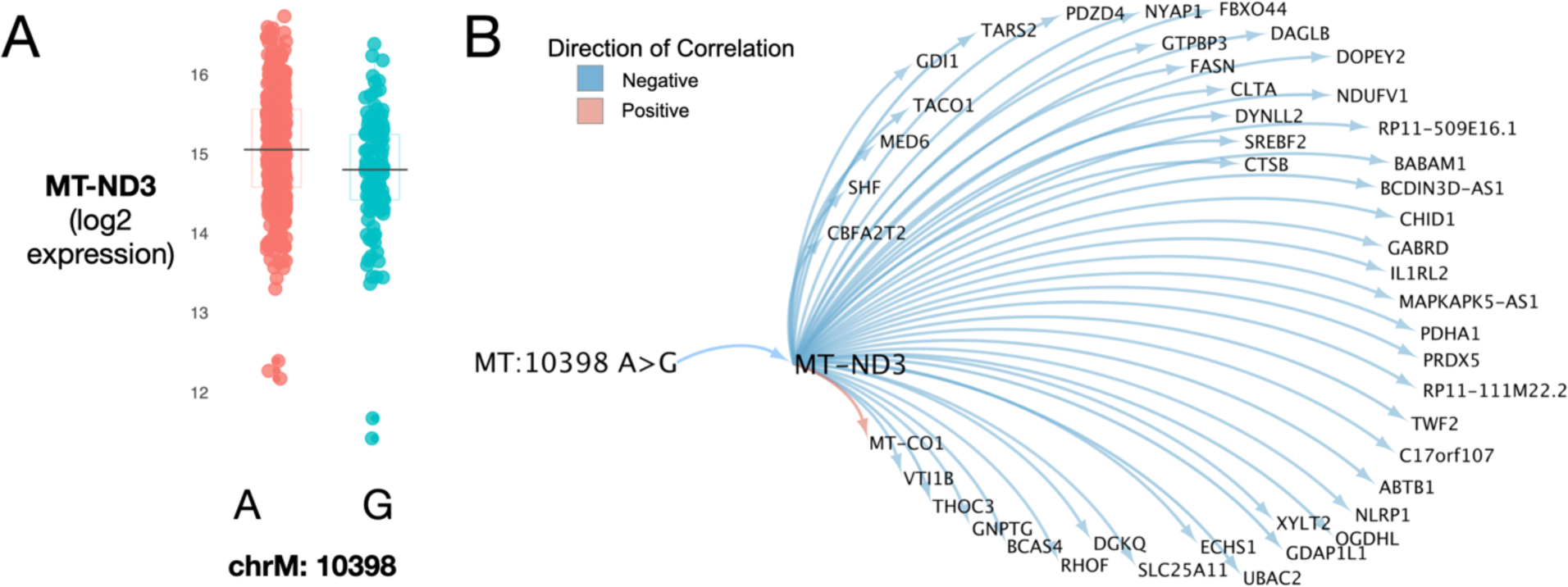
(A) Gene expression levels of MT-ND3 between dorsolateral prefrontal cortex samples with A or G allele for 10398 A>G. (B) Gene regulatory network constructed from neighbors downstream of MT-ND3, conditioned on MT:10398 A>G genotype. Top 45 strongest connections shown.

**Table 3.**
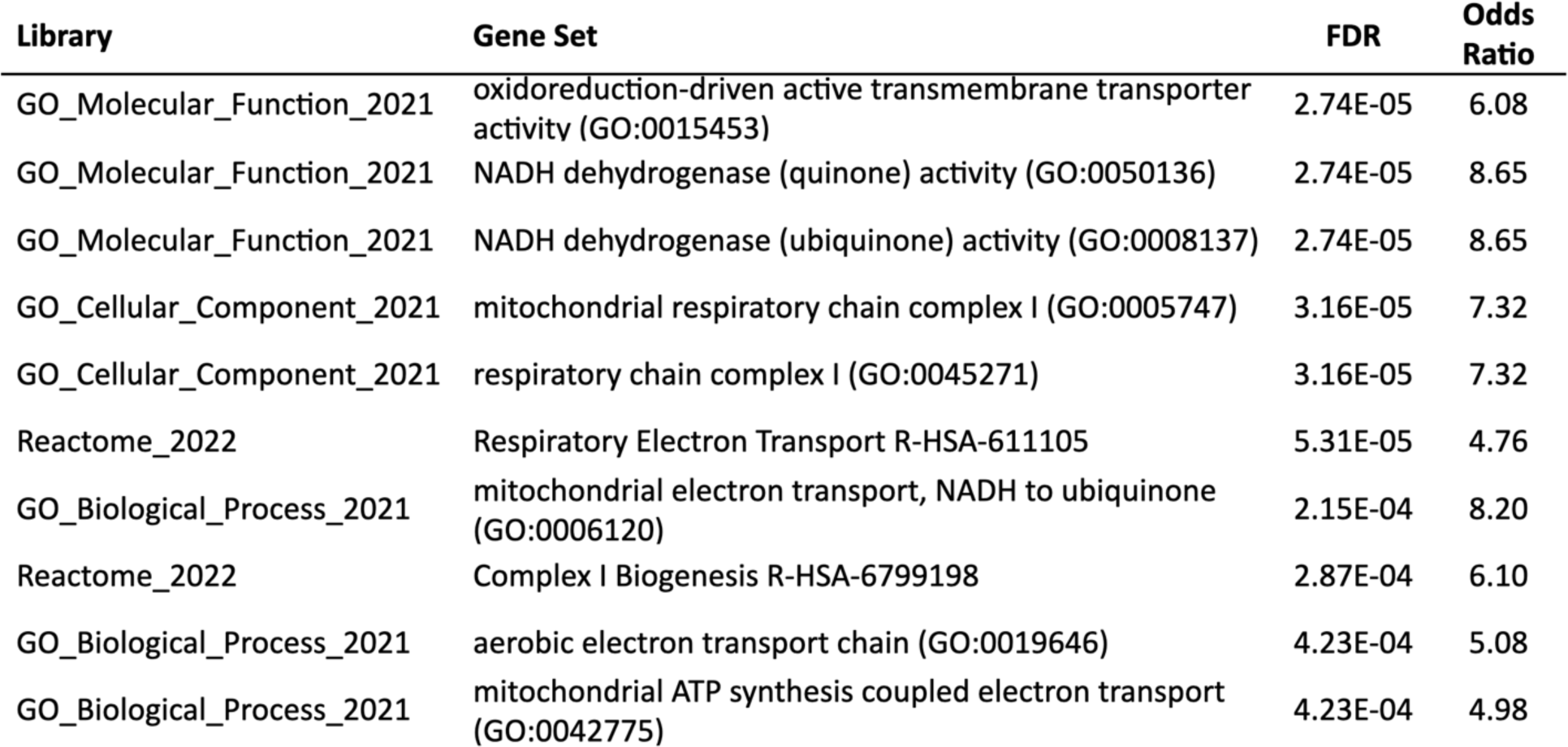
Selection of top Gene Ontology and Biological Pathways that are over-represented among the gene regulatory network constructed from neighbors downstream of MT-ND3, conditioned on MT:10398 A>G genotype.

| Library | Gene Set | FDR | Odds Ratio |
| --- | --- | --- | --- |
| GO_Molecular_Function_2021 | oxidoreduction-driven active transmembrane transporter activity (GO:0015453) | 2.74E-05 | 6.08 |
| GO_Molecular_Function_2021 | NADH dehydrogenase (quinone) activity (GO:0050136) | 2.74E-05 | 8.65 |
| GO_Molecular_Function_2021 | NADH dehydrogenase (ubiquinone) activity (GO:0008137) | 2.74E-05 | 8.65 |
| GO_Cellular_Component_2021 | mitochondrial respiratory chain complex I (GO:0005747) | 3.16E-05 | 7.32 |
| GO_Cellular_Component_2021 | respiratory chain complex I (GO:0045271) | 3.16E-05 | 7.32 |
| Reactome_2022 | Respiratory Electron Transport R-HSA-611105 | 5.31E-05 | 4.76 |
| GO_Biological_Process_2021 | mitochondrial electron transport, NADH to ubiquinone (GO:0006120) | 2.15E-04 | 8.20 |
| Reactome_2022 | Complex I Biogenesis R-HSA-6799198 | 2.87E-04 | 6.10 |
| GO_Biological_Process_2021 | aerobic electron transport chain (GO:0019646) | 4.23E-04 | 5.08 |
| GO_Biological_Process_2021 | mitochondrial ATP synthesis coupled electron transport (GO:0042775) | 4.23E-04 | 4.98 |

### The 10398A>G variant is significantly associated with MT-ND4 expression in LCLs

After performing RNA extraction and whole transcriptome RNA sequencing on all 60 LCL samples, we tested our data against previously discovered eQTLs in LCLs to determine if they were replicated within our samples. To do this, we integrated cis-eQTL data obtained from the GTEx portal for LCLs mapped in European-American subjects^51^. After filtering and pruning, we retained 764 SNPs that were also annotated as cis-eQTLs in GTEx and with available association statistics within our LCL data. We found an enrichment of GTEx cis-eQTL from our data in the same direction of effect (**Figure S1, Table S1**). Of the 764 SNPs, 98 of them were nominally significant from our data using a threshold of *P < 0.01*, whereas one would expect only 7.64 to be significant under the null hypothesis (*P = 7.4 x 10^-74^*). This result suggests that our LCL gene expression data is broadly concordant with the LCL data obtained from GTEx.

We then analyzed whether 10398A>G is associated with expression levels of any MT encoded genes. Of the 37 MT genes we examined for analysis, only 28 demonstrated robust expression in our data. We observed that 10398A>G is significantly associated with lower expression of MT-ND4 (*P=5.96 x 10^-9^, FDR=1.67 x 10^-7^*) (**Figure 4A, Table S2**). MT-ND4 (NADH Dehydrogenase 4), which is a different subunit from MT-ND3, is also part of the mitochondrial respiratory chain complex I. The G allele is also significantly associated with lower expression of MT-ATP8, MT-ND2, MT-ND4L, MT-ATP6 and MT-CYB (*FDR < 0.005*) (**Table S2**). Notably, there was no significant difference in expression detected for the MT-ND3 gene (as we had observed in the ROS/MAP DLPFC samples) despite the G allele causing a Thr114Ala missense mutation (*P=0.853, FDR=0.884*) (**Figure 4B**, **Table S2**).

**Figure 4.**
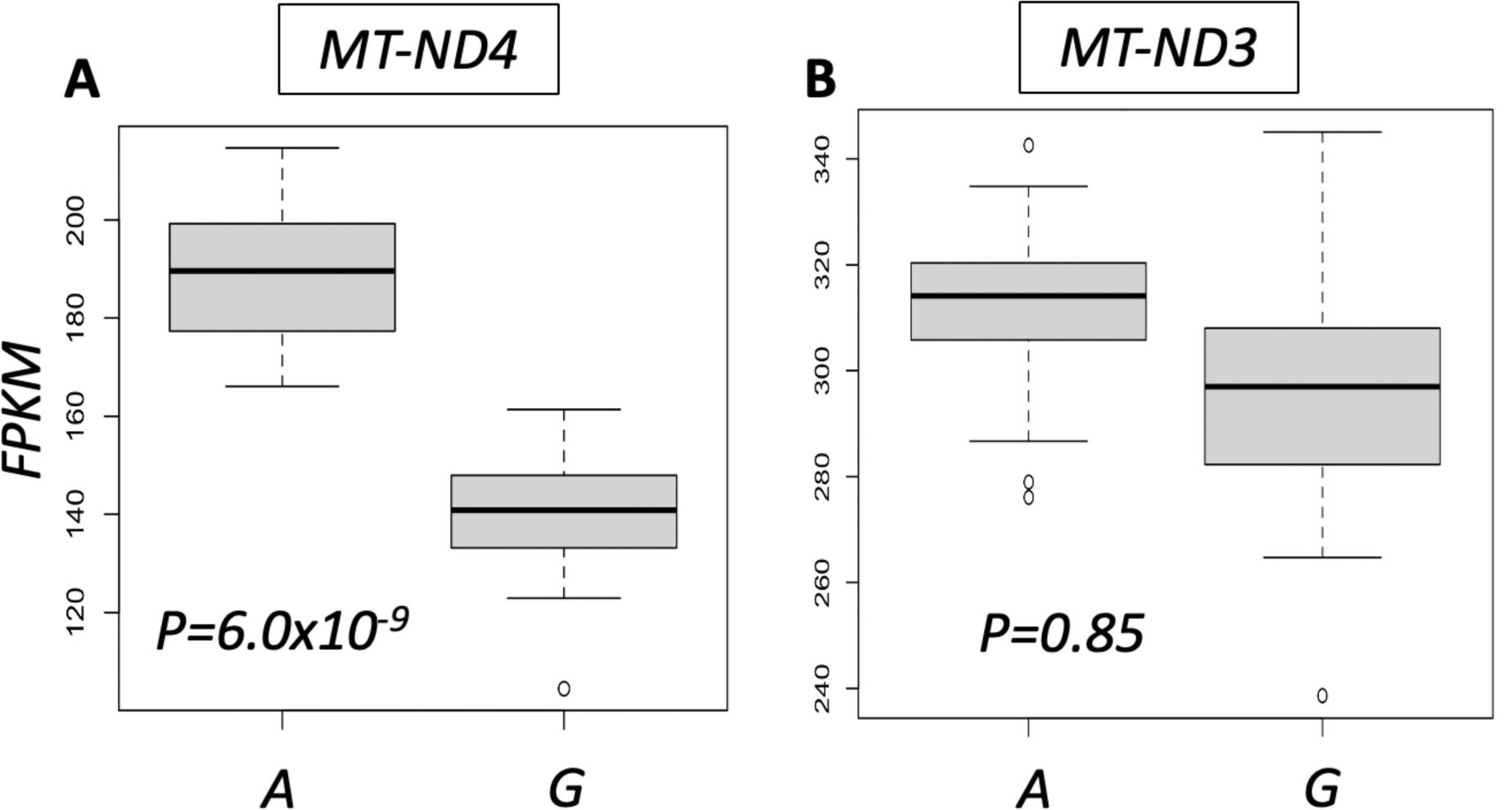
Comparing gene expression differences between LCL samples with the A allele versus the G allele of rs2853826. (A) Gene expression difference of MT-ND4. (B) Gene expression difference of MT-ND3.

### Testing for association of 10398A>G with mitochondrial DNA copy number in the LCLs

Given that the effect of 10398A>G on MT gene expression within the LCL samples was to uniformly decrease MT gene expression (**Table S2**), we hypothesized that this may reflect an effect on reducing overall MT copy number, as has been described for 10398A>G in a previous disease context^52^. We thus tested whether 10398A>G had any effect on MT copy number in the LCLs. We obtained genomic and mitochondrial DNA for each of the 60 PGP LCLs and performed multiplex PCR of a mitochondrial fragment, a fragment on the X-chromosome, and a fragment on chromosome 22 as a genomic control. The X-chromosome fragment serves as a positive control given that female samples should have significantly more X-chromosome DNA than male samples. Consistent with our expectation, the multiplex PCR approach demonstrated a significantly increased X-chromosome DNA copy number in our female samples compared to males, with clear separation between female and male samples (*P=1.28 x 10^-7^*) (**Figure 5A**). We performed the equivalent analysis to determine if 10398A>G is associated with altered MT copy number. The analysis did not detect any significant differences, suggesting that 10398A>G is not associated with altered MT copy number (*P=0.19*) (**Figure 5B**). In addition, we also performed MT copy number analysis using quantitative PCR and did not observe any significant difference with 10398A>G (*P=0.33*) (**Table S3**).

**Figure 5.**
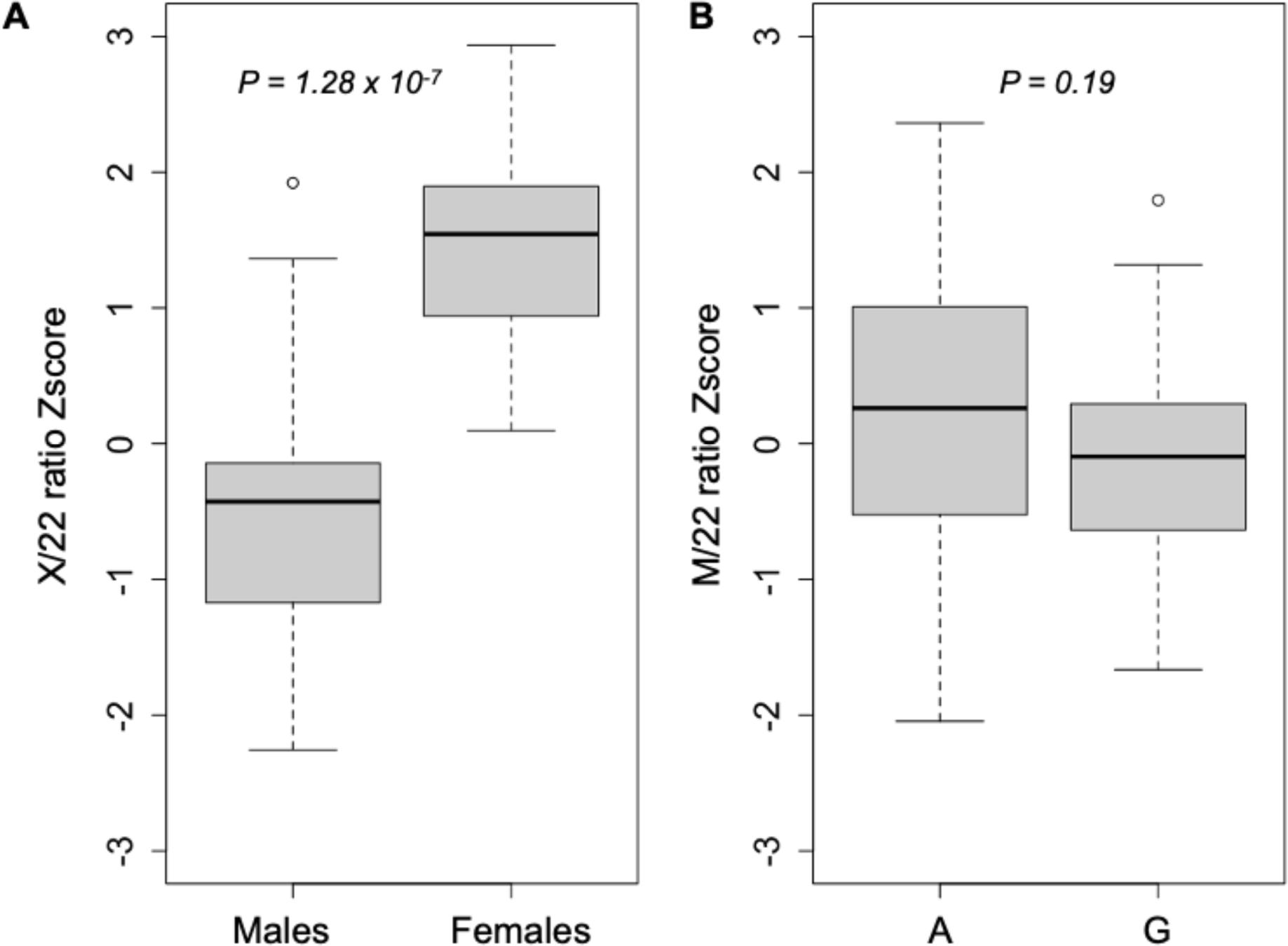
(A) DNA copy number analysis of the X-chromosome normalized against chromosome 22 between male and female samples. (B) Mitochondrial DNA copy number analysis normalized against chromosome 22 between the A and G allele at position 10398 of the mitochondrial DNA.

### 10398A>G is significantly associated with increased mitochondrial heteroplasmy

Finally, we quantified MT DNA heteroplasmy within the DNA obtained from our 60 PGP LCL samples by next-generation sequencing of the MT control region, where we had observed all MT heteroplasmy within brain derived samples. We designed primers to sequence neighboring regions chrM:16043-16238 and chrM:16469-262 of the MT DNA and calculated a heteroplasmy score for each site (See Materials and Methods). We used simple linear regression to look for association between age, sex, and MT copy number with individual heteroplasmy scores. There was no significant association between total heteroplasmy score and age (R^2^=-0.01609, *P*=0.79838), sex (R^2^=-0.01567, *P*=0.75984), or copy number mitochondrial Z-score (R^2^=-1.5x10^-3^, *P*=0.344). We then performed an association test of 10398A>G with the heteroplasmy scores and found that the minor G allele of 10398 was associated with significantly increased heteroplasmy at chrM:16469-262 (*P=0.011*) (**Figure 6B**). While there was a slight increase in heteroplasmy detected for chrM:16043-16238, the association statistic was not significant (*P=0.29*) (**Figure 6A**). Probing further into the individual sites, we found certain individual sites were contributing disproportionately to the overall increase in heteroplasmy (**Figure 7**).

**Figure 6.**
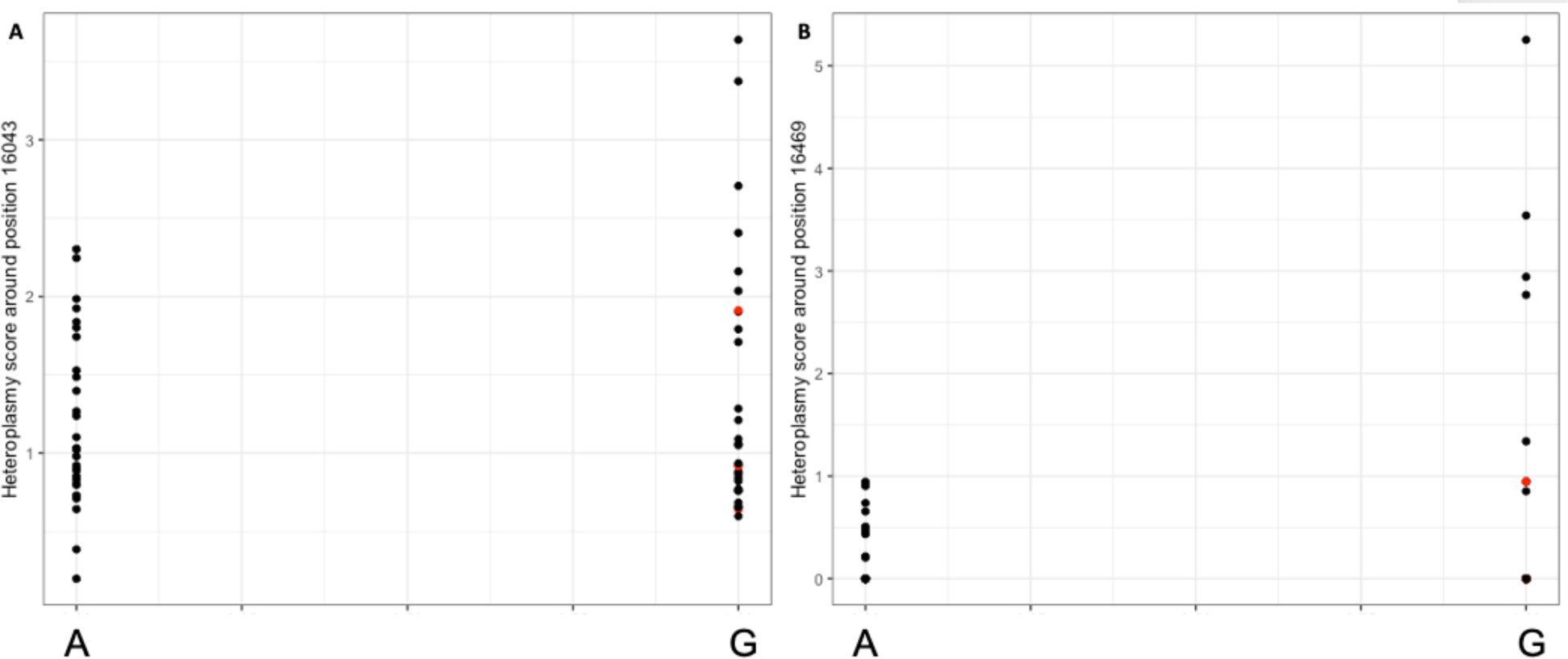
Association of A and G allele at position 10398 with (A) heteroplasmy score calculated for chrM:16043-16238 and (B) heteroplasmy score calculated for chrM:16469-262. Red dots indicate individuals of non-European ancestry stratified during PCA (See Materials and Methods).

**Figure 7.**
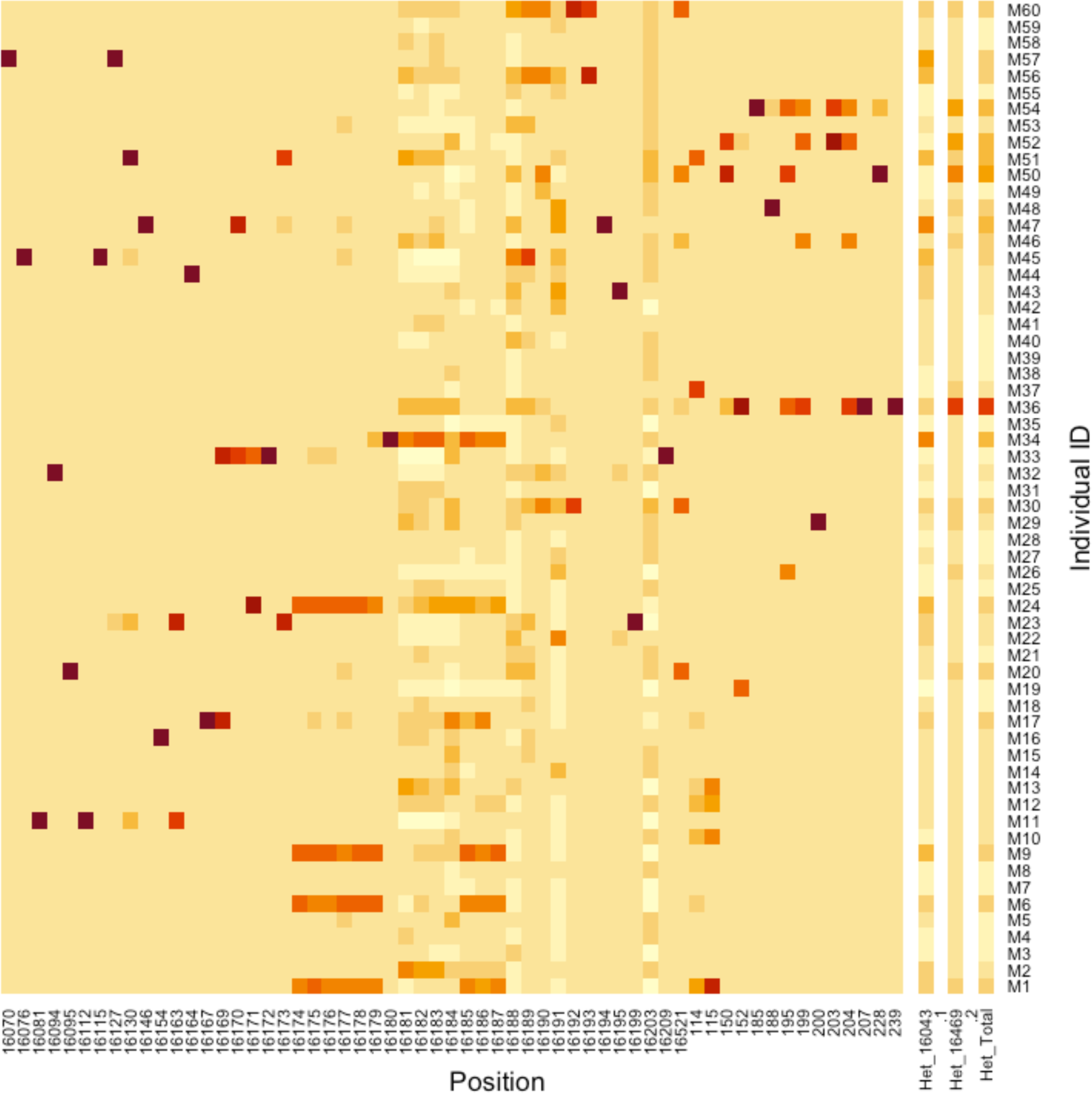
Individual heteroplasmic sites detected. Each row shows the heteroplasmy detected for each individual donor (M1 to M60). Darker regions show a higher heteroplasmy score for that site. The number on the x-axis shows selected positions on the mitochondrial DNA where heteroplasmy was detected. Het_16043 depicts the cumulative heteroplasmy for sites within chrM:16043-16238, Het_16469 depicts the cumulative heteroplasmy for sites within chrM:16469-262 and Het_Total depicts the combined heteroplasmy for both fragments.

We then performed an association test between all MT SNPs with overall heteroplasmy. We identified eight mitochondrial SNPs significantly associated (*P*<0.05) with total heteroplasmy score (**Table 4**). 10398A>G was significantly associated with higher heteroplasmy score (*P=*0.021). Furthermore, we computed the squared R^2^ correlation coefficient between each SNP identified as significantly associated with heteroplasmy score with 10398A>G and found that some of them were significantly correlated with 10398A>G (**Table 4**).

**Table 4.** List of mitochondrial SNPs that were associated with overall heteroplasmy.

| Position | rsID | Ref. | Alt. | MAF | Effect Size | SE | P-value | $R^2$ w. 10398 |
| --- | --- | --- | --- | --- | --- | --- | --- | --- |
| 73 | rs3087742 | G | A | 0.3333 | -0.671972 | 0.314507 | 0.0370148 | 0.486486 |
| 10398 | rs2853826 | A | G | 0.4737 | 0.720784 | 0.304438 | 0.0215864 | 1 |
| 12705 | rs2854122 | C | T | 0.2083 | 1.2496 | 0.394272 | 0.00247615 | 0.0362319 |
| 14766 | rs3135031 | T | C | 0.3667 | -0.675267 | 0.307857 | 0.0324439 | 0.565714 |
| 15043 | rs28357684 | G | A | 0.1167 | 1.94662 | 0.491097 | 0.000211222 | 0.0838574 |
| 15924 | rs2853510 | A | G | 0.15 | 1.15994 | 0.39716 | 0.00502691 | 0.208333 |
| 16129 | rs45134744 | G | A | 0.1583 | 1.27893 | 0.396306 | 0.0020912 | 0.00226131 |
| 16223 | rs2853513 | C | T | 0.2308 | 1.32282 | 0.415834 | 0.0025717 | 0.0737281 |

## Discussion

We analyzed large cohorts of DNA sequencing data to determine the extent of detectable mitochondrial heteroplasmy in various tissue types for different donor samples. We discovered that the 10398A>G (rs2853826) allele is significantly associated with mitochondrial heteroplasmy in the control region of brain samples. Given the reported associations between 10398A>G and a multitude of human diseases, we explored the functional consequence of 10398A>G with data available to us and discovered that it is associated with MT-ND3 expression levels in brain samples. We then characterized lymphoblastoid cell lines (LCLs) from the Harvard Personal Genome Project to determine the effect of 10398A>G in LCLs and discovered that it is associated with MT-ND4 expression levels and the MT-ND3 expression levels were not significantly altered in the LCLs. This suggests that 10398A>G could be affecting mitochondrial gene expression in a tissue or cell-type specific manner. Finally, from the sequencing data obtained from the LCLs, we found that 10398A>G is also associated with mitochondrial heteroplasmy, suggesting that heteroplasmy may be a general consequence of 10398A>G within multiple tissue types and potentially a mechanism by which 10398A>G confers an effect upon diverse human diseases.

Our initial motivation for exploring mitochondrial heteroplasmy was to determine its role in Alzheimer’s disease pathology. Early evidence indicated that post-mortem AD brains were enriched for heteroplasmic mitochondrial control region mutations^53^. Furthermore, recent evidence suggests mitochondrial DNA mutations can lead to mitochondrial dysfunction and increased Aβ deposition^54^. Our findings here lead us to the mitochondrial variant 10398A>G, which we found to be a strong predictor of heteroplasmic mutations in the mitochondrial control region. In an early study, 10398A>G was shown to be associated with AD risk^17^. However, a later study of a cohort of patients from Tuscany reported no mitochondrial haplotype group association with AD^55^. A later study in a cohort of Japanese individuals suggests little to no contribution of 10398A>G to AD^56^. Recently, there was a report of other mitochondrial variants associated with AD in a 254 individuals Tunisian cohort^57^. While that study did not find 10398A>G to be associated with AD, they identified other variants such as 5656A>G and 13759A>G to be marginally associated with AD. These studies suggest that the effect of 10398A>G on AD, if true, will be moderate and may not be observed in small cohort studies of only a few hundred patients. Nevertheless, it remains to be determined if the underlying cause of the association of 10398A>G to AD and other human diseases could be due to the effect of mitochondrial heteroplasmy on cellular function.

Besides AD, the role of mitochondrial heteroplasmy has also been studied for other neurological diseases. In autism spectrum disorders (ASD), researchers have discovered pathogenic heteroplasmic mitochondrial mutation being associated with ASD^58,59^. Perhaps the most common and well-studied mutation linked with mitochondrial heteroplasmy is 3243A>G, which has been associated with mitochondrial disorders and dysfunctions, including Mitochondrial Encephalopathy, Lactic Acidosis, Stroke-like episodes (MELAS), and Leigh syndrome^60^. As such, our finding of an inherited polymorphism 10398A>G being linked to mitochondrial heteroplasmy may inspire future research to investigate the role of mitochondrial heteroplasmy as a causal component of disease pathology.

Given the association of mitochondrial heteroplasmy with such a wide range of human diseases, and the previously reported progression from mitochondrial heteroplasmy to mitochondrial dysregulation and disruption of the OXPHOS pathway^61^, an exploration of the downstream effects of heteroplasmy and the 10398A>G variant on metabolic pathways may be warranted. Further evaluation of associations between the 10398A>G variant with other AD-associated phenotypes may prove to be fruitful. 10398A>G has previously been linked with Human Papillomavirus (HPV) positivity in the context of cervical cancer, indicating a potential modulation of host-virus interactions^52^. Given prior evidence of the role of Herpes simplex virus 1 (HSV-1) in AD pathogenesis in APOE-ε4 carriers^62–65^, one avenue of experimentation might be to test the viability of viral infection (e.g. HSV-1) in cell lines from individuals with the A versus G allele at 10398. This may provide better context for the relationships between the 10398A>G variant, mitochondrial heteroplasmy and damage, and AD pathology.

In summary, we have demonstrated the use of large scale whole-genome sequence data for analyzing mitochondrial heteroplasmy and performing cis-heteroplasmy-QTL analysis. Doing so led us to discover the association between 10398A>G and mitochondrial heteroplasmy. We then further characterize the effect of 10398A>G with the use of personalized LCL models. This highlights the utility of such resources as an effective way to perform *in-vitro* modeling for exploring genetic drivers of mitochondrial function in diseases, and suggests novel avenues for illuminating the role of heteroplasmy as a candidate mediating mechanism that links 10398A>G with diverse human diseases.

## Materials and Methods

### Whole genome sequences from the ROS, MAP, MSBB and Mayo cohorts

Mitochondrial variables (germline variation, heteroplasmy and copy number) were estimated on tissue samples with available whole genome sequences (WGS) generated by the Accelerating Medicines Partnership in Alzheimer’s Disease (AMP-AD). BAM files can be accessed at the AD Knowledge Portal (https://adknowledgeportal.org) for the MSBB (syn19987071), Mayo (syn19989379) and ROS/MAP (syn20068543) cohorts.

### Mitochondrial variant calling from brain derived whole genome sequences

Mitochondrial SNV and INDEL variants were called on existing WGS data using the gatk4 mitochondrial pipeline available here: https://github.com/gatk-workflows/gatk4-mitochondria-pipeline.

(1) Available BAM files were subset to reads with a primary mapping to MT genome using the gatk PrintReads function:

gatk PrintReads -R human_g1k_v37.fasta -L MT --read-filter

MateOnSameContigOrNoMappedMateReadFilter --read-filter

MateUnmappedAndUnmappedReadFilter -I Full.WGS.bam -O MT.bam

(2) MT.bam files were reheadered to exchange chromosomal notation from MT to chrM:

samtools view -H MT.bam | sed -e “s/SN:MT/SN:chrM/” | samtools reheader - MT.bam > chrM.bam

(3) chrM.bam files were indexed:

java -jar picard.jar BuildBamIndex I=chrM.bam

(4) chrM.bam files were submitted as inputs to Cromwell workflows designed for the gatk4-mitochondrial-pipeline

java -Dconfig.file=cromwell.singularity.conf -jar cromwell-47.jar run mitochondria-pipeline.remove.unpaired.wdl --inputs sample_config.json

(5) The called variants have been filtered by FilterMutectCalls to label false positives with a list of failed filters and true positives with PASS and only PASS variants were kept

gatk FilterMutectCalls -V output.vcf.gz --contamination-table contamination.table -O filtered.vcf.gz

gatk SelectVariants -R reference.fasta -V filtered.vcf.gz --exclude-filtered true -O final.vcf.gz

(6) Called variants are merged to a single file for each cohort, leaving uncalled sites missing: bcftools merge -l file.lst -o all.vcf

### Mitochondrial heteroplasmy quantitative trait loci analysis

Mitochondrial (MT) heteroplasmy quantitative trait loci (hetQTL) analysis was performed on whole genome sequences generated from dorsolateral prefrontal cortex (DLPFC) samples collected from 451 unique subjects within the Religious Orders Study (ROS) and Memory and Ageing Project (MAP). We applied a logistic regression approach, modelling the presence (or absence) of heteroplasmy at each MT site, as a function of each MT SNP genotype, while adjusting for AD status (NIA Reagan Score), Study (ROS or MAP), Sex, Age of death, Years of education, Post Mortem Interval, and Ancestry (estimated using the 10 largest population ancestry components precomputed from the WGS data). Within the ROS/MAP DLPFC samples, we identified 77 common MT variants (MAF >= 5%) and nine MT heteroplasmic sites that were detected in at least five subjects, which were carried forward into the hetQTL analysis. HetQTL associations were estimated using a generalized linear model using the R function “glm” and the family type “binomial”. Associations between each MT variant and MT heteroplasmic site were adjusted using the Benjamini-Hochberg approach for controlling the false discovery rate (FDR). Associations with an FDR < 0.05 were classified as significant. Results were annotated with symbols from overlapping genes, and separately with symbols from genes within 20 kilobases of MT variants and heteroplasmy sites using annotations from the GRCh38 transcriptome assembly.

### Mitochondrial expression quantitative trait loci analysis

Mitochondrial (MT) cis expression quantitative trait loci (cisQTL) analysis was performed across the set of samples with whole genome sequences (originating from any tissue) and RNA-sequences generated from dorsolateral prefrontal cortex (DLPFC) samples, comprising 573 unique subjects within the Religious Orders Study (ROS) and Memory and Ageing Project (MAP). Within the ROS/MAP DLPFC samples, we identified 212 common MT variants (MAF >= 5%). Transcriptomic data was subset to the 37 MT encoded genes. Genes were retained in the analysis if expression was detected in at least five samples, resulting in 23 included genes. Normalized abundance values were offset by a count of 1 and log2 transformed. Cis-eQTL analysis was performed using the R package, MatrixEQTL. Gene expression was modelled using a linear additive approach, including terms for MT variant genotype and while adjusting for AD status (NIA Reagan Score), Study (ROS or MAP), Sex, Age of death, Years of education, Post Mortem Interval, RNA Integrity Number (RIN), RNA-seq batch, and Ancestry (estimated using the 10 largest population ancestry components precomputed from the WGS data). Cis-eQTL relationships were defined as a maximum distance of 1 MB between a MT variant/gene pair, which was definitionally true for all examined MT pairs.

### Construction of MT-ND3 gene regulatory network

We performed causal inference testing^47^, to build a causal gene regulatory network focused on MT-ND3 in post-mortem DLPFC brain tissue collected as part of the ROS/MAP studies. This approach requires paired gene expression and genotype data for a large number of samples to establish the direction of regulation between MT-ND3 and its correlated genes.

Causal inference testing (CIT) has been well described previously^47^. Briefly, it offers a hypothesis test for whether a molecule (in this case, the expression of MT-ND3) is potentially mediating a causal association between a DNA locus (10398A>G), and some other quantitative trait (such as the expression of genes correlated with MT-ND3 and 10398A>G). Causal relationships can be inferred from a chain of mathematic conditions, requiring that for a given trio of loci (L), a potential causal mediator i.e., MT-ND3 (G) and a quantitative trait (T), the following conditions must be satisfied to establish that G is a causal mediator of the association between L and T:

a. L and G are associated
b. L and T are associated
c. L is associated with G, given T
d. L is independent of T, given G

We used the R software package “cit”^47^, to perform the causal inference test, calculating a false discovery rate using 1000 test permutations. Trios with a Qvalue < 0.05 were classified as significant, and the associated T genes were considered downstream of MT-ND3.

### Gene set enrichment testing

We then submitted the 1293 the downstream neighbors of MT-ND3 to the Enrichr webtool using the enrichR R package(23586463, 27141961, 33780170), specifically querying the “Reactome_2022”, “GO_Biological_Process_2021”, “GO_Cellular_Component_2021”, “GO_Molecular_Function_2021”, and “WikiPathway_2021_Human” gene set libraries. Enrichment results with an adjusted Pvalue < 0.05 were classified as significant.

### Personal Genome Project Lymphoblastoid Cell Lines and Whole Genome Sequencing Data

The lymphoblastoid cell lines (LCL) from the Personal Genome Project (PGP) cohort were obtained from the Coriell Institute for Medical Research (https://www.coriell.org/). The whole genome sequencing data (WGS) for PGP samples were obtained from the PGP website (https://pgp.med.harvard.edu). There were a total of 123 unique donors with both LCLs and WGS data available. We performed data mining of the 123 donors and determined that 30 donors carry the rs2853826 “G” allele of which 3 individuals also have the “GCT” allele (**Table S4**). Most of these individuals were self-reported as having European ancestry except for 4 donors (2 Chinese, 1 African-American and 1 Hispanic). We then selected an additional 30 donor samples carrying the rs2853826 “A” allele as control samples bringing the total number of donors analyzed to 60. We also performed sanger sequencing of the rs2853826 locus for all 60 samples to confirm their rs2853826 genotype (**Figure S2**).

### LCL cell culture, DNA and RNA extraction and RNA sequencing

Lymphoblastoid cell lines (LCL) were cultured in RPMI medium supplemented with 10% Fetal Bovine Serum (FBS) and 1% penicillin-streptomycin. These cultured LCLs were maintained at 37 °C. DNA from the LCLs were extracted using the AccuPrep® Genomic DNA Extraction Kit (Bioneer). RNA was extracted from the LCLs using the PureLink™ RNA Mini Kit (Thermofisher). The extracted RNA for whole transcriptome sequencing was sent to Psomagen for sequencing. Preparation of samples for RNA-Seq analysis was performed using the TruSeq RNA Sample Preparation Kit (Illumina, San Diego, CA). Briefly, rRNA was depleted from total RNA using the Ribo-Zero rRNA Removal Kit (Human/Mouse/Rat) (Illumina, San Diego, CA) to enrich for coding RNA and long non-coding RNA, following the TruSeq Stranded Total RNA Sample Prep Guide, Part # 15031048 Rev. The Ribo-Zero libraries were sequenced on the Illumina NovaSeq 6000 System with 151 nucleotide paired end reads, according to the standard manufacturer’s protocol (Illumina, San Diego, CA). The raw sequence reads were aligned to human genome hg38 (ensembl_GRCh38, GenBank Assembly ID GCA_000001405.15) with the star aligner (v2.5.2b) and gene level expression (read counts) were summarized by the “--quantMode GeneCounts” parameter in star.

### Testing mitochondrial DNA copy number

We used a multiplex PCR approach to ascertain mitochondrial DNA copy number. We designed 3 primer pairs to target the mitochondrial DNA, a region on the X-chromosome and a region on chromosome 22. The primers for the mitochondrial DNA were TACACATGCAAGCATCCCCG for the forward primer and ATCACTGCTGTTTCCCGTGG for the reverse primer. These primers target chrM:692-826 which results in a 135bp PCR fragment. The primers for the X-chromosome DNA were ATCCCCGTGTGGTAGTCTCC for the foward primer and AGTTGCCAGACGTCTTAAAGTCC for the reverse primer. These primers target chrX:13086693-13086892 which results in a 200bp PCR fragment. The primers for chromosome 22 were CAGAGGCTCAGAGAGGTCATCT for the forward primer and CCTAAGGTTGAGTTTGGTCTCCC for the reverse primer. These primers target chr22:37103514-37103839 which results in a 326bp PCR fragment. 50ng of genomic and mitochondrial DNA was used as template DNA for the multiplex PCR reaction. All primer sequences are given in the 5’ to 3’ orientation. The genome coordinates were provided using the GRCh37 (hg19) assembly of the human genome. The resulting PCR product was sent for analysis using a 5200 Fragment Analyzer System (Agilent) and the quantification for each PCR fragment is given for each sample. The multiplex PCR was performed in 2 separate batches. Batch one had samples M1-15, M31-45 and batch 2 had samples M16-30, M46-60. We then calculated the X/22 and M/22 ratios for each sample and obtained a Zscore for both statistics by normalizing against each batch’s sample mean and standard deviation (**Table S5**).

### PCR amplification and sequencing of the mitochondrial control region

We PCR amplified 2 regions of the mitochondrial DNA for sequencing to determine mitochondrial heteroplasmy. For the first site, chrM:16043-16238, we used CTTTCCCTACACGACGCTCTTCCGATCTNNNNNNgatttgggtaccacccaagt as the forward primer and GGAGTTCAGACGTGTGCTCTTCCGATCTgttgaaggttgattgctgtact as the reverse primer. As for the second site, chrM:16469-262, which consist of a 365 bp fragment, we used CTTTCCCTACACGACGCTCTTCCGATCTNNNNNNcttgggggtagctaaagtga as the forward primer and GGAGTTCAGACGTGTGCTCTTCCGATCTggctgtgcagacattcaatt as the reverse primer. For these primer sequences, the lower-case bases are homologous to the target region while the upper-case bases are overhangs required for the second round PCR. The amplified fragments then undergo a second round of PCR where the illumina indexes and primer sequences are attached using i7 and i5 primers. The i7 primer sequence was CAAGCAGAAGACGGCATACGAGAT[i7]GTGACTGGAGTTCAGACGTGTGCTCTTCCGATCT and the i5 primer sequence was AATGATACGGCGACCACCGAGATCTACAC[i5]ACACTCTTTCCCTACACGACGCTCTTCCGA TCT. The i7 and i5 sequence consist of a unique 8 bp sequence to identify the dual-indexed illumina sequence library. All sequences are depicted in the 5’ to 3’ orientation. The primer sequences were synthesized using Custom DNA Oligos Synthesis Services (Thermofisher). We used a Q5 High-Fidelity DNA Polymerase (NEB) and ran the PCR on a ProFlex Thermal Cycler (Thermofisher).

The DNA libraries were then pooled and sent for whole-genome sequencing (Psomagen). We performed Illumina paired-end sequencing (read 150 bases) on the HiSeq X Ten system, with different index barcodes for each sample resulting in roughly one million paired-end reads per sample on average (T**able S6**).

### Calculating heteroplasmy score

We performed alignment of the PCR-amplified sequencing reads for each sample to the hg19 assembly of the human genome using bwa^66^. We mapped the aligned sequences to a fasta file containing the primers and mitochondrial genome sequences using the mpileup function of samtools^67^. From the mpileup file, we constructed a statistic to measure the degree of heteroplasmy across our nine sites of interest and the surrounding bases using the following method.

For each of the 60 individuals in our cohort, we computed the fraction of non-reference alleles to the total number of reads at all PCR-amplified positions (each mtDNA site between bases 16043 to 16238 and bases 16469 to 262). We first identified any sites with measurable heteroplasmy (noted in the formula below with the indicator function) which are sites with allele fraction estimates between 0.05 and 0.95 (**Table S7**). Sites with non-reference allele fraction less than or equal to 0.05 or greater than or equal to 0.95 are assigned as stably inherited SNPs, and the heteroplasmy measure at these sites are set to zero. We then weighted each heteroplasmic site by the allele fraction and summed them up to generate heteroplasmy scores for each individual.

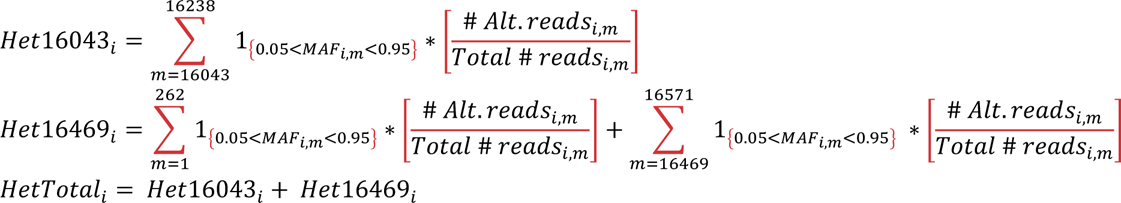

Het16043_i_ is the heteroplasmy score of individual *i* generated from sites between bases 16043 to 16238. Het16469_i_ is the heteroplasmy score of individual *i* generated from sites between bases 16469 to 262 and HetTotal_i_ is the heteroplasmy score of individual *i* generated by taking the sum of Het16043_i_ and Het16469_i_.

### Quality control and association analysis of MT SNPs with heteroplasmy

Plink 1.90b6.9^68^ was used to perform quality control and association analysis. We ensured that no individuals had a genotyping rate less than 20%, discordant sex data, or were duplicated or shared significant identity-by-descent. Because of limited sample size, SNPs with minor allele frequency less than or equal to 0.09 as well as those not in Hardy-Weinburg equilibrium (*P<1.0x10^-6^*) were removed. Of the 60 samples in our cohort of LCLs, we found that three individuals of non-European ancestry (2 Chinese, 1 African-American) were stratified in PCA. (**Figure S3**). We used Plink to implement a generalized linear regression model for the remaining 42 MT SNPs for the 60 LCL samples in our cohort, including the first two principal components from PCA as covariates, with HetTotal_i_ and calculated the overall association statistics (**Table S8**).

## Acknowledgements

The results published here are in whole or in part based on data obtained from the AD Knowledge Portal (https://adknowledgeportal.org).

Data pertaining to the ROS and MAP studies were provided by the Rush Alzheimer’s Disease Center, Rush University Medical Center, Chicago. Data collection was supported through funding by NIA grants P30AG10161 (ROS), R01AG15819 (ROSMAP; genomics and RNAseq), R01AG17917 (MAP), R01AG30146, R01AG36042 (5hC methylation, ATACseq), RC2AG036547 (H3K9Ac), R01AG36836 (RNAseq), R01AG48015 (monocyte RNAseq) RF1AG57473 (single nucleus RNAseq), U01AG32984 (genomic and whole exome sequencing), U01AG46152 (ROSMAP AMP-AD, targeted proteomics), U01AG46161(TMT proteomics), U01AG61356 (whole genome sequencing, targeted proteomics, ROSMAP AMP-AD), the Illinois Department of Public Health (ROSMAP), and the Translational Genomics Research Institute (genomic). Additional phenotypic data can be requested at www.radc.rush.edu. Data pertaining to the MSBB study was generated from postmortem brain tissue collected through the Mount Sinai VA Medical Center Brain Bank and was provided by Dr. Eric Schadt from Mount Sinai School of Medicine.

The Mayo study data was led by Dr. Nilüfer Ertekin-Taner, Mayo Clinic, Jacksonville, FL as part of the multi-PI U01 AG046139 (MPIs Golde, Ertekin-Taner, Younkin, Price). Samples were provided from the following sources: The Mayo Clinic Brain Bank. Data collection was supported through funding by NIA grants P50 AG016574, R01 AG032990, U01 AG046139, R01 AG018023, U01 AG006576, U01 AG006786, R01 AG025711, R01 AG017216, R01 AG003949, NINDS grant R01 NS080820, CurePSP Foundation, and support from Mayo Foundation. Study data includes samples collected through the Sun Health Research Institute Brain and Body Donation Program of Sun City, Arizona. The Brain and Body Donation Program is supported by the National Institute of Neurological Disorders and Stroke (U24 NS072026 National Brain and Tissue Resource for Parkinsons Disease and Related Disorders), the National Institute on Aging (P30 AG19610 Arizona Alzheimers Disease Core Center), the Arizona Department of Health Services (contract 211002, Arizona Alzheimers Research Center), the Arizona Biomedical Research Commission (contracts 4001, 0011, 05-901 and 1001 to the Arizona Parkinson’s Disease Consortium) and the Michael J. Fox Foundation for Parkinsons Research.

BR is supported in part by NIA grants U01AG061835 and R21AG063068. QW is supported in part by NIA grants U01AG061835. ETL and YC are supported in part by NIA grant U01AG061835. BR and DM are grateful for funding support received from the Arizona Alzheimer’s Consortium.

## Ethics Declaration

### Ethics approval

The use of the human cell lines for research in this work was determined to not be human subjects research by the IRB of UMass Chan Medical School (H00021419).

