## Supplementary Figures for "Modeling of mitochondrial genetic polymorphisms reveals induction of heteroplasmy by pleiotropic disease locus MT:10398A>G"

### Figure S1

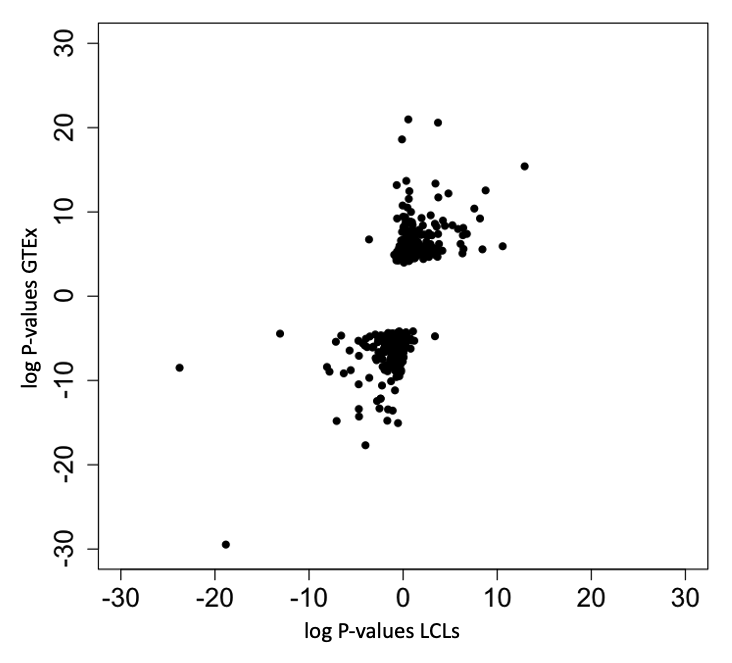

Comparing the log P-values of gene expression QTLs between the data obtained from our LCL samples versus LCL data obtained from GTEx. Negative log P-values indicate genes annotated as having lower gene expression with the effect allele whereas positive log P-values indicate genes annotated as having higher gene expression with the effect allele.

### Figure S2

Sanger sequencing results for all 60 samples to verify the mitochondria allele status for each sample. The fragment sequence depicted for each sample is ACTGA**[A/G]**CCGAATT.

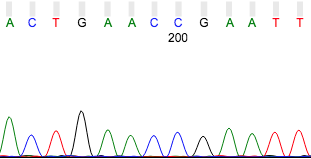

M1-MT1_FOR3 (Allele detected: A)

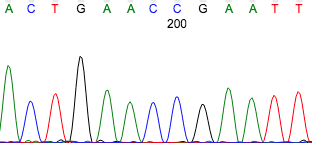

M2-MT1_FOR3 (Allele detected: A)

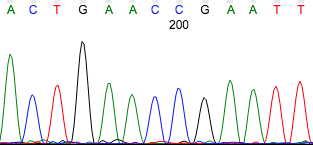

M3-MT1_FOR3 (Allele detected: A)

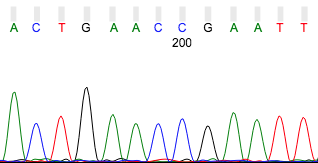

M4-MT1_FOR3 (Allele detected: A)

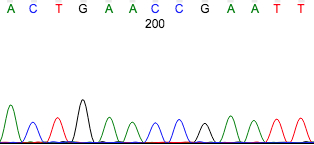

M5-MT1_FOR3 (Allele detected: A)

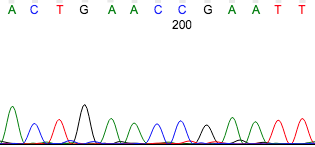

M6-MT1_FOR3 (Allele detected: A)

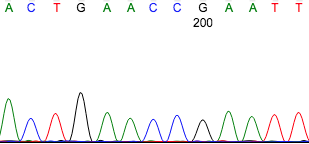

M7-MT1_FOR3 (Allele detected: A)

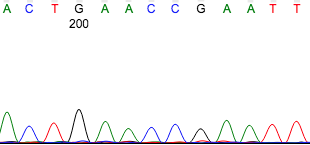

M8-MT1_FOR3 (Allele detected: A)

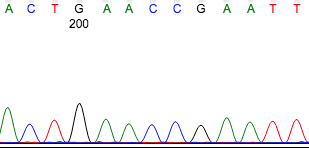

M9-MT1_FOR3 (Allele detected: A)

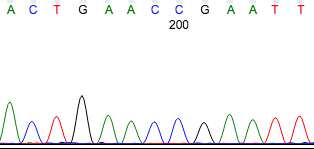

M10-MT1_FOR3 (Allele detected: A)

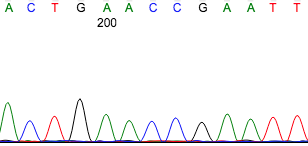

M11-MT1_FOR3 (Allele detected: A)

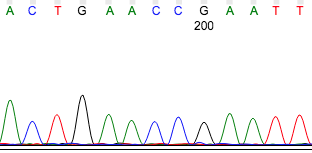

M12-MT1_FOR3 (Allele detected: A)

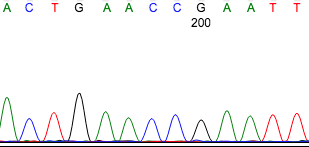

M13-MT1_FOR3 (Allele detected: A)

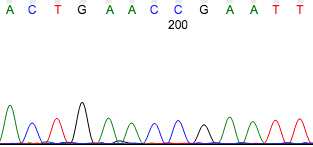

M14-MT1_FOR3 (Allele detected: A)

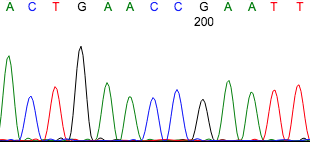

M15-MT1_FOR3 (Allele detected: A)

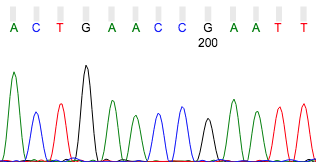

M16-MT1_FOR3 (Allele detected: A)

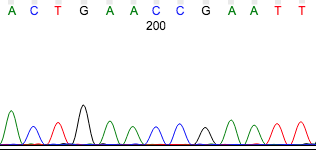

M17-MT1_FOR3 (Allele detected: A)

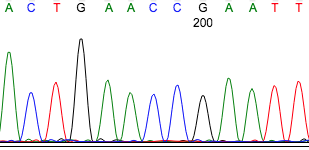

M18-MT1_FOR3 (Allele detected: A)

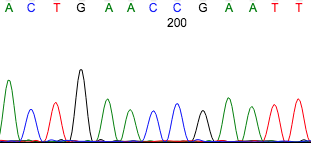

M19-MT1_FOR3 (Allele detected: A)

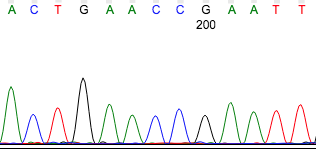

M20-MT1_FOR3 (Allele detected: A)

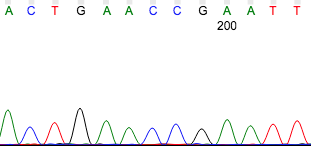

M21-MT1_FOR3 (Allele detected: A)

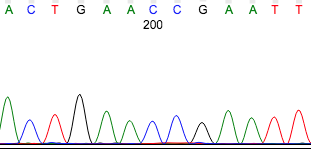

M22-MT1_FOR3_R (Allele detected: A)

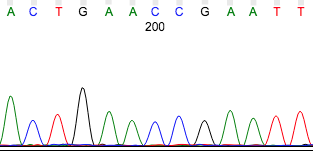

M23-MT1_FOR3 (Allele detected: A)

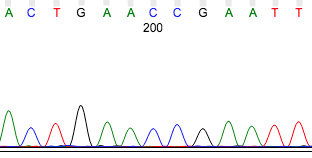

M24-MT1_FOR3 (Allele detected: A)

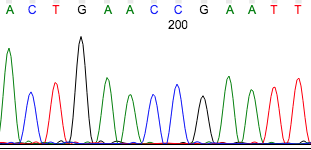

M25-MT1_FOR3 (Allele detected: A)

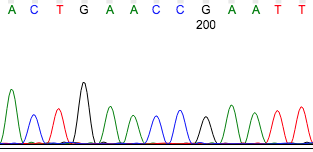

M26-MT1_FOR3 (Allele detected: A)

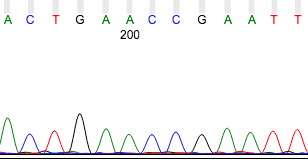

M27-MT1_FOR3 (Allele detected: A)

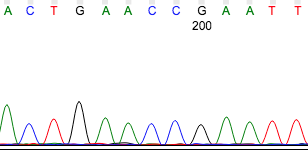

M28-MT1_FOR3 (Allele detected: A)

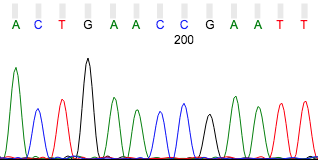

M29-MT1_FOR3 (Allele detected: A)

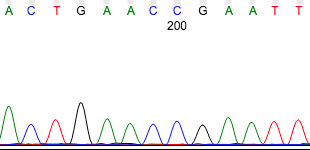

M30-MT1_FOR3 (Allele detected: A)

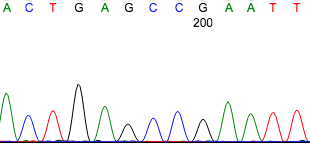

M31-MT1_FOR3 (Allele detected: G)

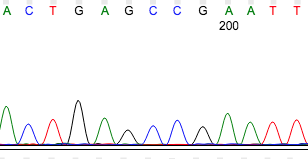

M32-MT1_FOR3 (Allele detected: G)

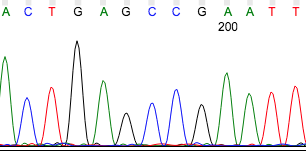

M33-MT1_FOR3 (Allele detected: G)

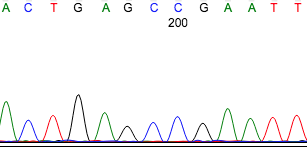

M34-MT1_FOR3 (Allele detected: G)

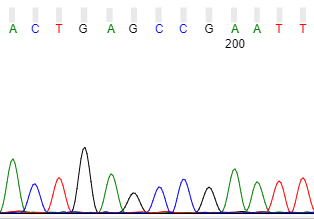

M35-MT1_FOR3 (Allele detected: G)

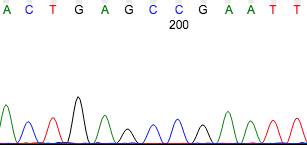

M36-MT1_FOR3 (Allele detected: G)

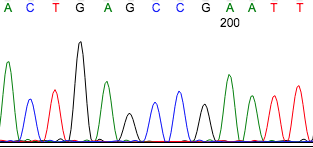

M37-MT1_FOR3 (Allele detected: G)

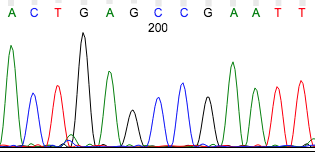

M38-MT1_FOR3 (Allele detected: G)

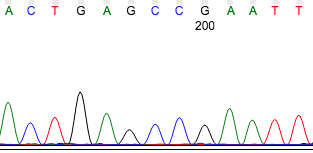

M39-MT1_FOR3 (Allele detected: G)

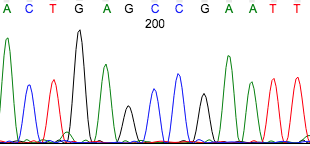

M40-MT1_FOR3 (Allele detected: G)

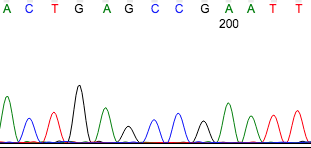

M41-MT1_FOR3 (Allele detected: G)

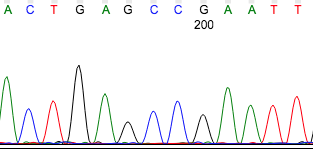

M42-MT1_FOR3 (Allele detected: G)

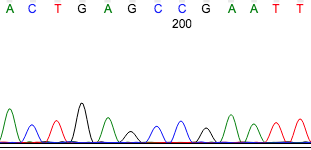

M43-MT1_FOR3 (Allele detected: G)

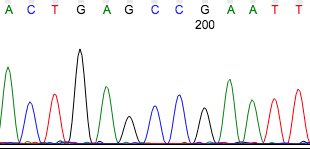

M44-MT1_FOR3 (Allele detected: G)

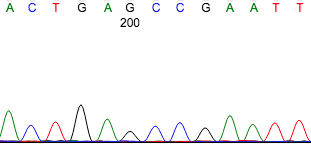

M45-MT1_FOR3 (Allele detected: G)

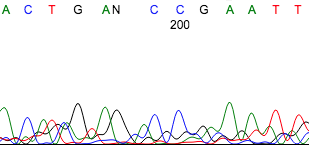

M46-MT1_FOR3 (Allele detected: G, Poor quality)

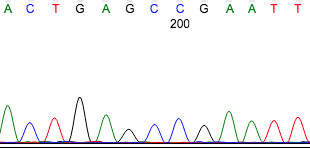

M47-MT1_FOR3 (Allele detected: G)

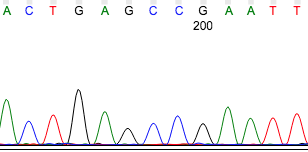

M48-MT1_FOR3 (Allele detected: G)

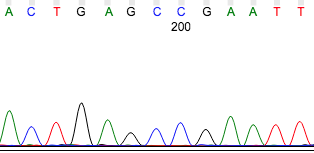

M49-MT1_FOR3 (Allele detected: G)

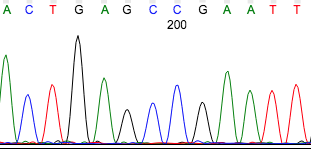

M50-MT1_FOR3 (Allele detected: G)

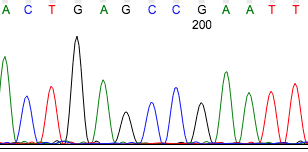

M51-MT1_FOR3 (Allele detected: G)

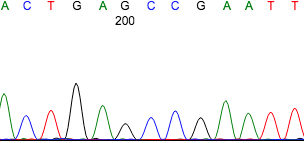

M52-MT1_FOR3 (Allele detected: G)

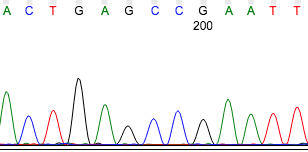

M53-MT1_FOR3 (Allele detected: G)

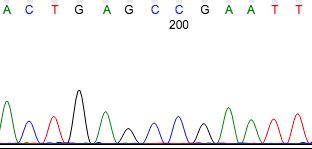

M54-MT1_FOR3 (Allele detected: G)

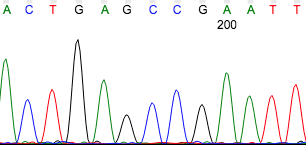

M55-MT1_FOR3 (Allele detected: G)

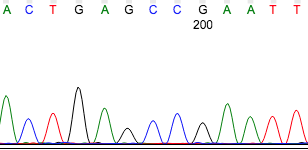

M56-MT1_FOR3 (Allele detected: G)

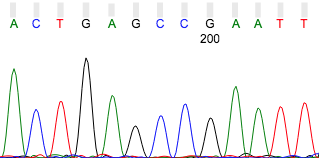

M57-MT1_FOR3 (Allele detected: G)

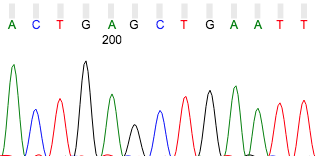

M58-MT1_FOR3 (Allele detected: GCT)

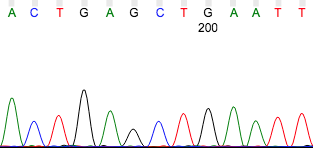

M59-MT1_FOR3 (Allele detected: GCT)

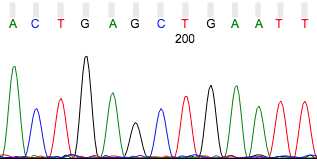

M60-MT1_FOR3 (Allele detected: GCT)

#### Figure S3

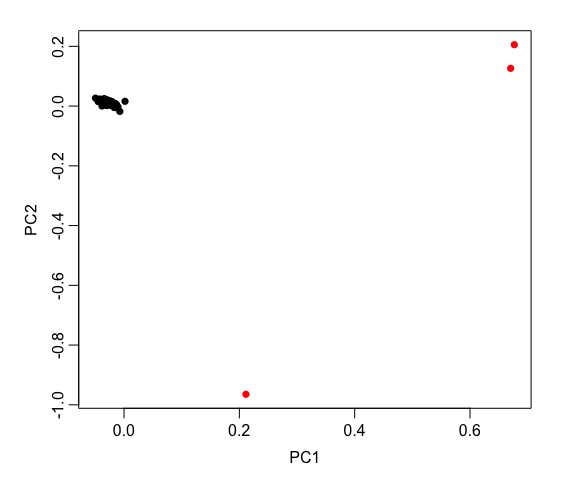

Principal component analysis using the genomic SNPs extracted for each sample. The first 2 principal components (PC1 and PC2) are depicted. The outlier samples are indicated in red.
